# Differential regulation of flower transpiration during abiotic stress in plants

**DOI:** 10.1101/2021.11.29.470467

**Authors:** Ranjita Sinha, Sara I. Zandalinas, Yosef Fichman, Sidharth Sen, Aurelio Gómez-Cadenas, Trupti Joshi, Felix B. Fritschi, Ron Mittler

**Affiliations:** Division of Plant Sciences and Technology, College of Agriculture Food and Natural Resources and Interdisciplinary Plant Group. University of Missouri. Columbia, MO 65211, USA; Departamento de Ciencias Agrarias y del Medio Natural, Universitat Jaume I, Castelló de la Plana, 12071, Spain; Institute for Data Science and Informatics and Interdisciplinary Plant Group, University of Missouri, Columbia, MO 65211, USA; Department of Health Management and Informatics, and Christopher S. Bond Life Sciences Center, University of Missouri, Columbia, MO 65211, USA; Department of Surgery, University of Missouri School of Medicine, Christopher S. Bond Life Sciences Center, University of Missouri, Columbia, MO 65201, USA

**Keywords:** Climate change, crop, drought, flower, global warming, heat stress, soybean, stomata, stress combination, tobacco, transpiration, water deficit, yield

## Abstract

Heat waves, occurring during droughts, can have a devastating impact on yield, especially if they happen during the flowering and seed set stages of the crop cycle. Global warming and climate change are driving an alarming increase in the frequency and intensity of combined drought and heat stress episodes, critically threatening global food security. Previous studies revealed that during a combination of drought and heat stress stomata on leaves of many plants are closed, preventing cooling by transpiration. Because high temperature is detrimental to reproductive processes, essential for plant yield, we measured the inner temperature, transpiration, and sepal stomatal aperture of closed soybean flowers, developing on plants subjected to a combination of drought and heat stress. Here, we report that during a combination of drought and heat stress soybean plants prioritize transpiration through flowers over transpiration through leaves by opening their flower stomata, while keeping their leaf stomata closed. This acclimation strategy, termed ‘differential transpiration’, lowers flower inner temperature by about 2-3°C, protecting reproductive processes at the expense of vegetative tissues. Manipulating stomatal regulation, stomatal size and/or stomatal density of flowers could therefore serve as a viable strategy to enhance the yield of different crops and mitigate some of the current and future impacts of global warming and climate change on agriculture.

**One sentence summary:** During stress conditions that result in higher flower inner temperature plants use a differential transpiration strategy to protect reproductive processes at the expense of vegetative tissues.

## INTRODUCTION

The unyielding increase in atmospheric and oceanic temperatures, termed ‘global warming’, is causing drastic changes in our climate, termed ‘climate change’ (Lobell et al., 2011; Steg, 2018; Bailey-Serres et al., 2019; Alizadeh et al., 2020; Overpeck and Udall, 2020; von der Gathen et al., 2021; Zandalinas et al., 2021; Zhai et al., 2021). As a result, large areas of our planet are increasingly exposed to floods or extended droughts combined with extreme temperatures (Mazdiyasni and AghaKouchak, 2015; Alizadeh et al., 2020; Overpeck and Udall, 2020; Rivero et al., 2021; Zandalinas et al., 2021). Historically, extended droughts combined with heat waves have been the cause of catastrophic reductions in agricultural productivity estimated at billions of dollars per episode (*e.g.,* the drought and heat wave combination events occurring during the summers of 1980 and 1988 in the US resulted in losses to agriculture estimated at 33 and 44 billion dollars, respectively; https://www.ncdc.noaa.gov/billions/; Mittler, 2006; Lobell et al., 2011; Rivero et al., 2021). Because global warming and climate change are increasing the frequency and intensity of drought and heat stress combination events worldwide, more studies are needed to understand how crops and other plants respond to this type of stress combination (Mazdiyasni and AghaKouchak, 2015; Alizadeh et al., 2020; Rivero et al., 2021; Zandalinas et al., 2021; Zhai et al., 2021). Agricultural experience, as well as multiple studies conducted with different crops, revealed that the effects of drought and heat stress combination on yield of many major grain crops is most severe when the stress combination occurs during the reproductive stage of plant growth (Rollins et al., 2013; Mahrookashani et al., 2017; Lawas et al., 2018; Liu et al., 2020; Cohen et al., 2021b; Rivero et al., 2021; Sinha et al., 2021).

Plant reproduction, *i.e.,* the developmental process of flower organs (including stamens and stigma), the maturation of pollen and egg cells, pollen shedding, interactions with stigma, germination, growth and eventually fertilization, as well as embryo development and seed filling, are all highly sensitive to elevated temperatures (Santiago and Sharkey, 2019; Djanaguiraman et al., 2020; Chaturvedi et al., 2021; Sze et al., 2021). It was recently proposed that the tightly synchronized nature of the developmental programs involved in these processes, as well as their reliance on certain stress-related programs (*e.g.,* the stress-associated desiccation program of pollen grains), reactive oxygen species (ROS) and hormone signaling, under non-stress conditions, makes them more sensitive to stress (Sinha et al., 2021; Sze et al., 2021). Stresses such as drought or heat, or their combinations, could therefore disrupt these tightly synchronized processes by triggering the activation of stress programs, and/or the accumulation of different hormones, ROS, or other signals, out of sync with the proper developmental process, leading to the production of immature or malnourished pollen, egg cell programmed cell death, and other disruptive processes that decrease yield (Martin et al., 2013; Lassig et al., 2014; Daneva et al., 2016; Zhang et al., 2020; Sinha et al., 2021; Sze et al., 2021).

Many important grain crops such as wheat (*Triticum aestivum*), rice (*Oryza sativa*) and soybean (*Glycine max*) are self-pollinating and do not require vectors such as insects or wind for cross-pollination (Liu et al., 2006). In many legumes and important grass species, self-pollination occurs even before the flower opens (*i.e.,* the pollen is transferred to the stigma of the same flower within the closed flower; termed cleistogamy or pseudocleistogamy; Campbell et al., 1983; Takahashi et al., 2001). Although under controlled non-stressed conditions cleistogamy/pseudocleistogamy protects many aspects of the reproductive process from external factors such as low humidity, UV radiation, pathogens, and/or other potential stressors, under conditions of drought, or drought combined with heat stress, when transpiration is suppressed, the internal temperature of the flower could rise to high levels that would inhibit reproduction (Lawas et al., 2018; Wei et al., 2018; Sinha et al., 2021).

Transpiration in plants is primarily controlled by changes in stomatal aperture and the water vapor pressure differential between the plant and the atmosphere (Will et al., 2013; Lawson and Matthews, 2020). When stomata are open transpiration can occur at a higher rate and cool the plant. This was demonstrated for leaves of different plants subjected to heat stress (Zhou et al., 2015; Balfagón et al., 2019; Zandalinas et al., 2020a). In contrast, during drought stomata are closed to prevent water loss and plant temperature increases due to lack of transpiration. Interestingly, during a combination of drought and heat stress, stomata on leaves of many different plant species are closed and leaf temperature is higher than that of plants subjected to heat alone (Rizhsky et al., 2002; Rizhsky et al., 2004; Carmo-Silva et al., 2012; Zandalinas et al., 2016; Cohen et al., 2021a). Because the temperature of reproductive processes (occurring within the flowers of cleistogamous plants) plays such a critical role in the overall yield of many crops, we studied how a combination of drought and heat stress (that has a devastating impact on yield, Mittler, 2006; Lobell et al., 2011; Rollins et al., 2013; Mahrookashani et al., 2017; Lawas et al., 2018; Liu et al., 2020; Cohen et al., 2021b; Rivero et al., 2021; Sinha et al., 2021), would impact flower stomatal aperture, transpiration, and inner temperature in two different plants: soybean and tobacco (*Nicotiana tabacum*). Here, we report that during a combination of drought and heat stress, plants prioritize transpiration through flowers over transpiration through leaves by opening their flower stomata, while keeping their leaf stomata closed. This strategy, termed ‘differential transpiration’, lowers flower temperature by about 2-3°C, and represents a newly discovered acclimation mechanism of plants to different abiotic stresses that result in higher inner flower temperature (*e.g.,* combinations of drought, pathogen infection, mechanical injury, high CO_2_, or air pollution, such as ozone, that cause stomatal closure, with heat stress). Manipulating stomatal regulation, stomata size, and/or stomata number (*i.e.,* stomatal density) of flowers could therefore serve as a viable strategy to enhance the yield of different crops in the face of our uncertain current and future climates.

## RESULTS

### Leaf and flower temperature of plants subjected to a combination of water deficit and heat stress

To induce conditions of water deficit (WD), heat stress (HS) and a combination of WD and HS (WD+HS), we grew soybean plants (*Glycine max*, *cv Magellan*) in controlled growth chambers. When plants began to flower (R1 stage) we induced conditions of WD, HS and WD+HS (Cohen et al., 2021a) and maintained these conditions for 10 days before starting to analyze and sample leaves and flowers. Using this design, we made sure that the new leaves and flowers we studied (R2 stage) developed under the different stress conditions. As shown in Figure 1A, as well as reported previously for different plant species (Rizhsky et al., 2002; Rizhsky et al., 2004; Carmo-Silva et al., 2012; Zandalinas et al., 2016; Cohen et al., 2021a), compared to plants subjected to WD or HS alone, leaf temperature of plants subjected to WD+HS was higher. To determine whether flowers of plants subjected to WD+HS exhibit a similar higher temperature (compared to flowers of plants subjected to HS or WD), we measured the internal temperature of flowers using a thermocouple thermometer probe (Figure 1B). For this analysis we used soybean flowers at stage II and III (unopen flowers undergoing self-pollination; Supplemental Figure S1) from plants grown under conditions of WD, HS, WD+HS, and control (CT) (Figure 1B). As shown in Figure 1C, the inner flower temperature of flowers that developed under conditions of WD+HS combination was higher than that of flowers grown under conditions of CT, HS or WD alone. Water potential (Ψ; psi, measured in MPa) is a measure of water pressure and osmotic potential and is typically low in tissues subjected to WD, HS, or WD+HS, potentially indicating water loss and tissue desiccation (Sattar et al., 2020; Cohen et al., 2021a). In addition to increased temperature (Figures 1A and C), the water potential of leaves (Figure 1D) and flowers (Figure 1E) from plants subjected to WD+HS was lower compared to that of leaves and flowers grown under conditions of CT, HS or WD alone. Taken together, the results presented in Figure 1 demonstrate that flowers of plants subjected to WD+HS have a high internal temperature that is coupled with low water potential.

**Figure 1.**
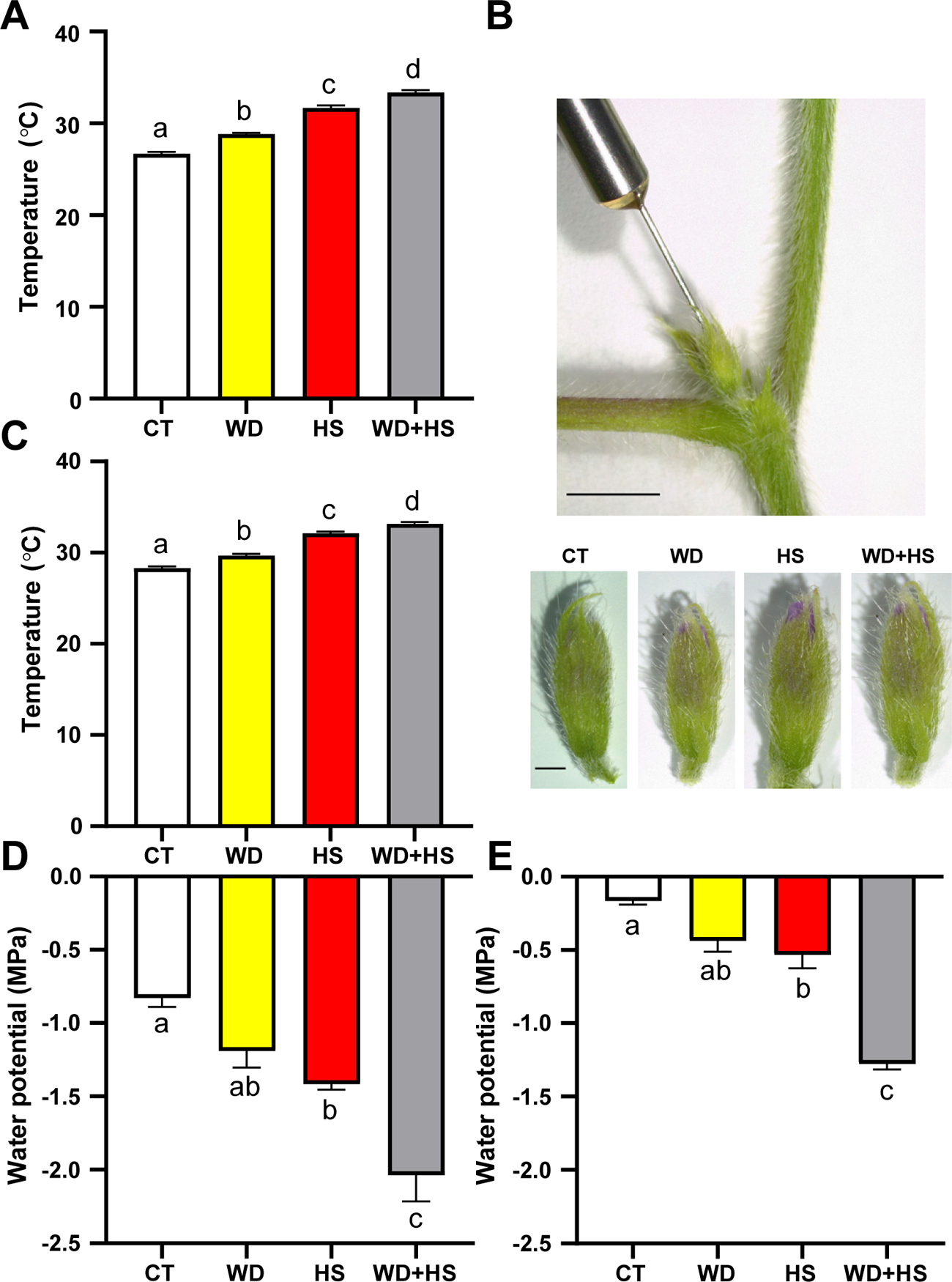
Leaf and flower temperature of soybean plants subjected to a combination of water deficit and heat stress. **A,** Leaf temperature of soybean plants subjected to control (CT), heat stress (HS), water deficit (WD), or WD+HS. **B,** Representative image of the experimental set up used to measure soybean inner flower temperature with a thermocouple thermometer probe (top; Bar is 5 mm) and representative images of closed (stage II and III; Supplemental Figure S1) soybean flowers developing under the different stress treatments (bottom; Bar is 1 mm). **C,** Inner flower temperature of soybean flowers from plants subjected to CT, WD, HS, or WD+HS conditions. **D,** Water potential (Ψ; psi, measured in MPa) of soybean leaves subjected to CT, WD, HS, or WD+HS. **E,** Water potential of soybean flowers from plants subjected to CT, WD, HS, or WD+HS. All experiments were repeated at least three times with at least 10 plants per biological repeat. Data are presented as mean ± SE. Different letters denote significance at P < 0.05 (ANOVA followed by a Tukey’s post hoc test). Abbreviations: CT, control; HS, heat stress; MPa, mega pascal; WD, water deficit.

### Stomatal aperture and transpiration of flowers and leaves from plants subjected to a combination of water deficit and heat stress

Stomatal aperture, stomatal conductance, and transpiration are key physiological parameters that determine plant temperature and water potential (Nilson and Assmann, 2007; Hsu et al., 2021; Lawson and Matthews, 2020). We therefore measured these parameters in leaves and flowers of plants subjected to WD+HS. In agreement with our previous findings obtained with soybean, tobacco, and Arabidopsis (Rizhsky et al., 2002; Rizhsky et al., 2004; Zandalinas et al., 2016; Cohen et al., 2021a), leaf stomatal aperture, stomatal conductance and transpiration remained high in plants subjected to HS, decreased in plants subjected to WD, and decreased in plants subjected to WD+HS (Figures 2A-C). These findings suggest that in contrast to HS, leaves subjected to WD+HS could not be cooled via transpiration (Rizhsky et al., 2002; Rizhsky et al., 2004; Mittler, 2006; Carmo-Silva et al., 2012; Zandalinas et al., 2016; Cohen et al., 2021a), and experience higher temperatures (Figure 1A). In contrast to leaves, flower (sepal) stomatal aperture, and whole flower stomatal conductance and transpiration, were high in plants subjected to WD+HS, as well as in plants subjected to HS, and low in plants subjected to WD (Figures 2D and E). This finding suggests that during a combination of WD+HS stomata of flowers (sepals) respond differently than stomata of leaves and remain open enabling cooling via transpiration. Interestingly, the inner temperature of flowers subjected to WD+HS was high (Figure 1C), despite the ongoing transpiration (Figure 2F). This observation could be explained by differences in the thickness of flowers and leaves. While soybean flower buds have a diameter of about 1.5-2 mm (Figure 1B) soybean leaves are much thinner (about 0.12-0.15 mm) and can be cooled by transpiration much more easily.

**Figure 2.**
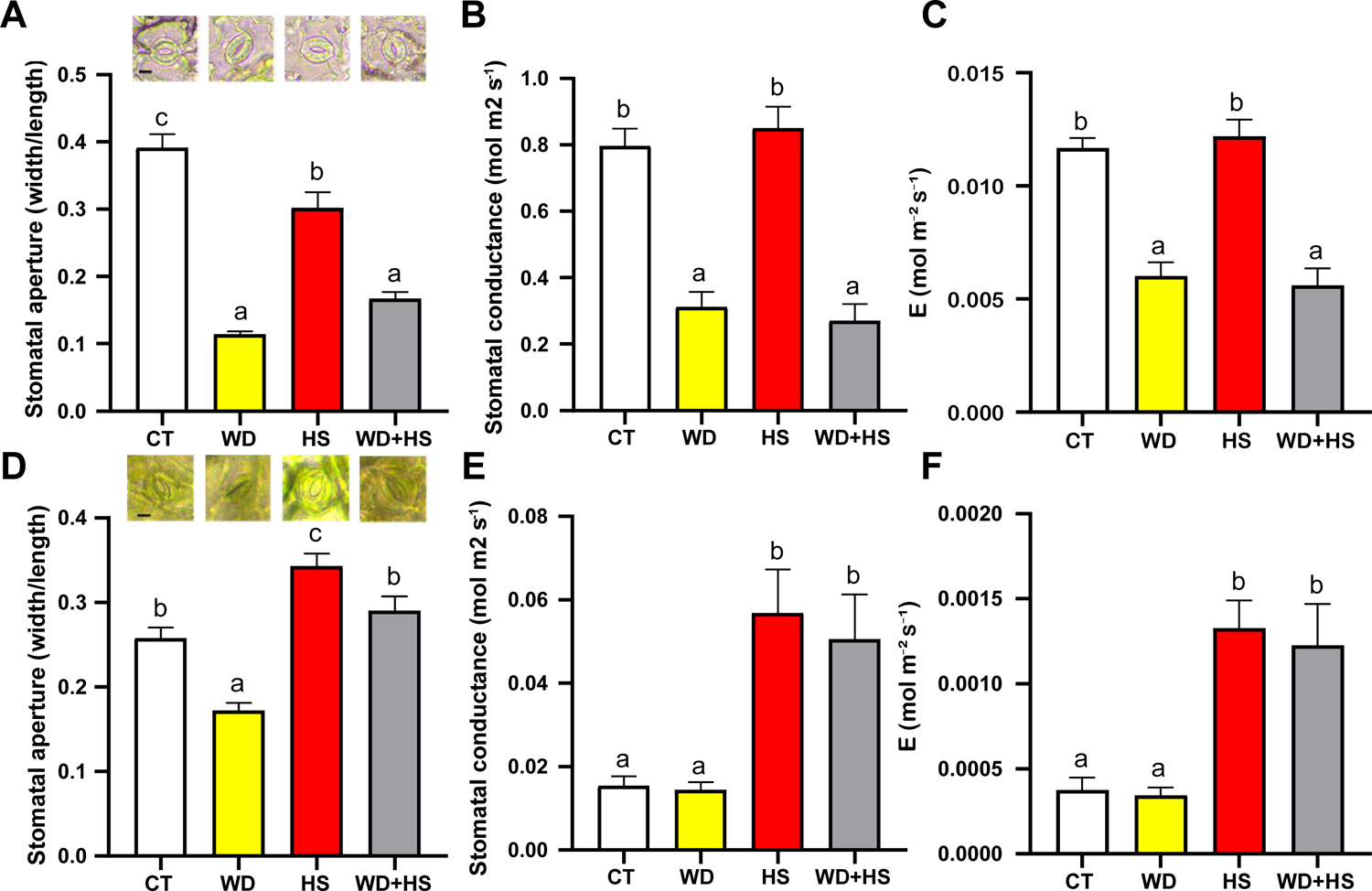
Stomatal aperture and transpiration of flowers and leaves from plants subjected to a combination of water deficit and heat stress. **A-C,** Stomatal aperture (**A**), stomatal conductance (**B**), and transpiration (**C**) of soybean leaves from plants subjected to control (CT), heat stress (HS), water deficit (WD), or WD+HS. **D-E.** Stomatal aperture (**D**), stomatal conductance (**E**), and transpiration (**F**) of soybean flowers from plants subjected to CT, HS, WD, or WD+HS. All experiments were repeated at least three times with at least 10 plants per biological repeat. At least 400 stomata from each treatment were measured. Data are presented as mean ± SE. Different letters denote significance at P < 0.05 (ANOVA followed by a Tukey’s post hoc test). Representative images of stomata are shown in **A** and **D**. Bar is 10 μm. Abbreviations: CT, control; E, transpiration; HS, heat stress; WD, water deficit.

### Stomatal density of leaves and flowers developed under conditions of water deficit and heat stress combination

Plants display a high degree of plasticity when grown under diverse environmental conditions (Chater et al., 2014; Zhu, 2016; Caine et al., 2019; Sakoda et al., 2019; Lloyd and Lister, 2021; Markham and Greenham, 2021). Among the different phenotypes plants can display in response to different growth conditions is a change in the density (number per area) of stomata appearing on the surface of newly developing leaves (Chater et al., 2014; Caine et al., 2019; Sakoda et al., 2019). The differential responses of stomata from sepals and leaves during WD+HS, as well as the lower levels of transpiration measured from whole flowers compared to leaves (Figure 2), prompted us to examine whether the number of stomata forming on these organs (*i.e.,* stomatal density), during their development under the stress conditions applied in our study, would also be different. As shown in Figure 3A, the stomatal density of leaves developed under HS or WD+HS was higher than that of leaves grown under WD or CT conditions, with stomatal density of leaves from plants grown under WD+HS being the highest. In contrast, as shown in Figure 3B, the stomatal density of sepals from plants subjected to HS was not significantly different from that of plants grown under CT or WD, while the stomatal density of sepals developing under WD+HS was higher than all other treatments. Although the overall density of stomata on sepals was lower than that on leaves (Figure 3), the average size of stomata found on sepals was larger than that of leaves (Figure 2; Supplemental Figure S2). The results presented in Figures 2 and 3 suggest that the developmental response of leaves and flowers (sepals) to WD+HS (*i.e.,* increase in stomatal density; Figure 3) is similar, while the physiological response of these two different organs (*i.e.,* opening or closing of stomatal aperture; Figure 2) is very different.

**Figure 3.**
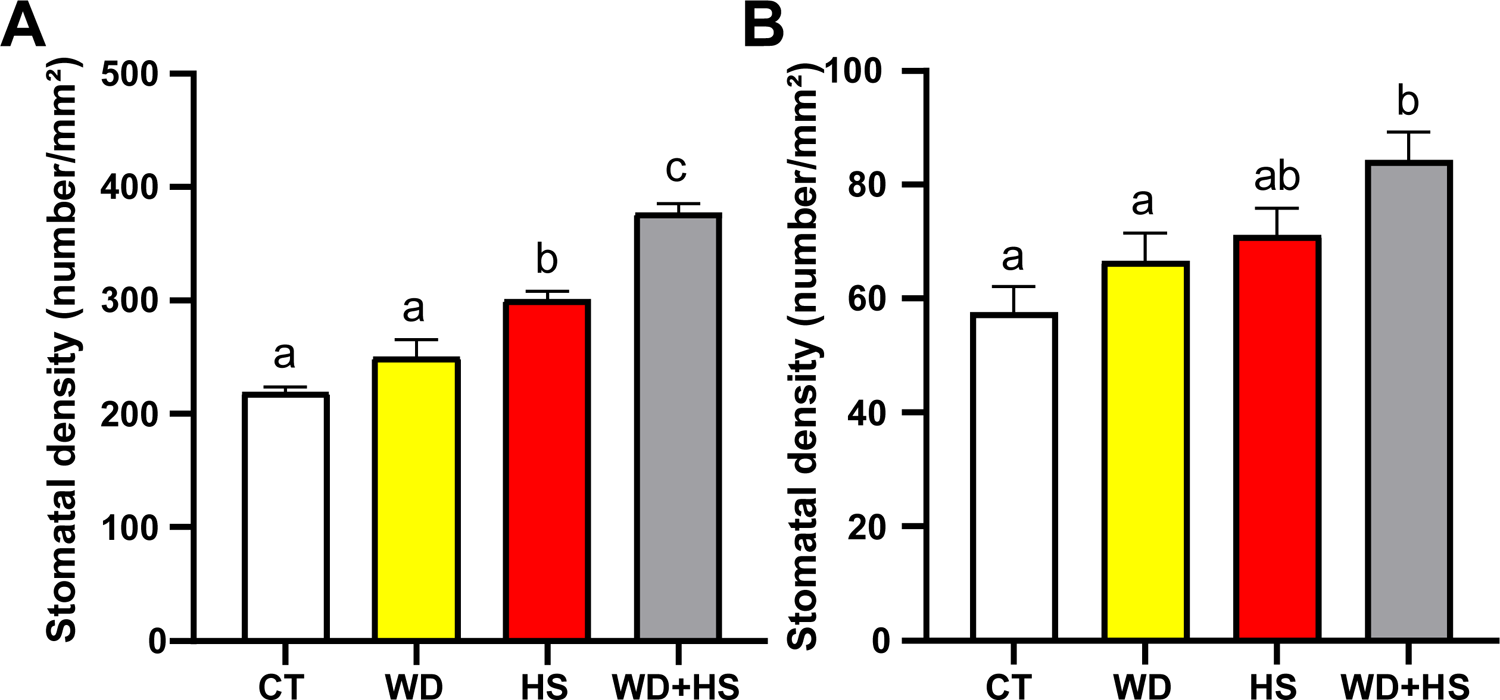
Stomatal density of leaves and flowers developed under conditions of water deficit and heat stress combination. **A**, stomatal density of leaves from plants subjected to control (CT), heat stress (HS), water deficit (WD), or WD+HS. **B**, Stomatal density of sepals from plants subjected to CT, HS, WD, or WD+HS. All experiments were repeated at least three times with at least 10 plants per biological repeat. At least 100 microscopic fields were counted from each treatment. Data are presented as mean ± SE. Different letters denote significance at P < 0.05 (ANOVA followed by a Tukey’s post hoc test). Abbreviations: CT, control; HS, heat stress; WD, water deficit.

### External application of ABA to flowers results in stomatal closure, and sealing of stomata results in elevated flower temperature under conditions of water deficit and heat stress combination

Stomatal aperture, conductance and overall transpiration are regulated in plants by various signals (Nilson and Assmann, 2007; Buckley, 2019; Hsu et al., 2021). Among these, abscisic acid (ABA) is well known to play a key role in triggering stomatal closure (Nilson and Assmann, 2007; Lozano-Juste and Cutler, 2016; Buckley, 2019; Hsu et al., 2021; Zhang et al., 2021). Because stomata of flowers from plants subjected to WD+HS combination were open, while stomata of leaves from the same plants were closed (Figure 2), we tested whether external application of ABA would cause stomatal closure in flowers (sepals) from plants subjected to the stress combination. As shown in Figure 4A, application of ABA to flowers grown under CT, HS, or WD+HS resulted in stomatal closure. In contrast, application of ABA to flowers from plants grown under WD did not change the stomatal aperture, as these stomata were already closed. The results presented in Figure 4A suggest that stomata of flowers subjected to WD+HS did not become resistance to external ABA application.

**Figure 4.**
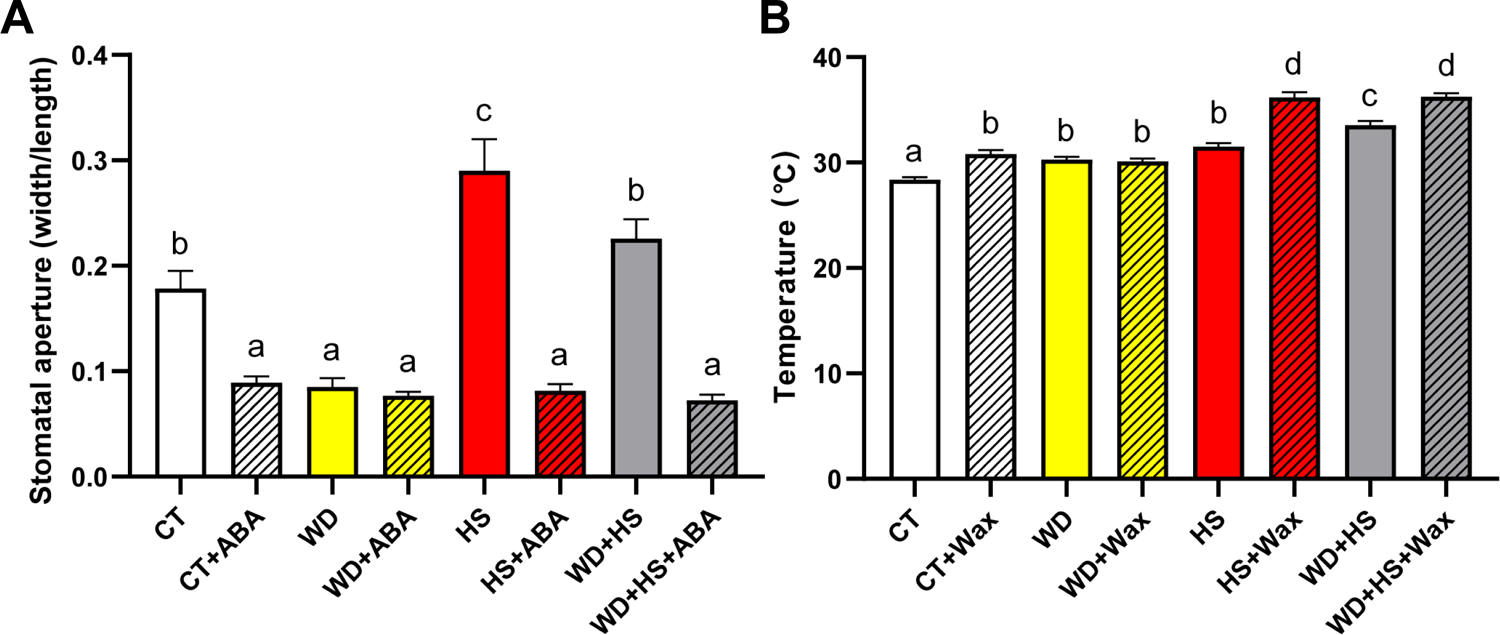
External application of abscisic acid (ABA) to flowers results in stomatal closure and sealing of stomata results in elevated flower temperature under conditions of water deficit and heat stress combination. **A**, Stomatal aperture of sepals from plants subjected to control (CT), heat stress (HS), water deficit (WD), or WD+HS, 60 min following application of water or ABA. **B,** Inner flower temperature of flowers from plants subjected to CT, WD, HS, or WD+HS, coated or uncoated with a thin layer of wax for 120 min. All experiments were repeated at least three times with at least 10 plants per biological repeat. Data are presented as mean ± SE. Different letters denote significance at P < 0.05 (ANOVA followed by a Tukey’s post hoc test). Abbreviations: ABA, abscisic acid; CT, control; HS, heat stress; WD, water deficit.

A possible reason to why flowers would keep their sepal stomata open, maintaining transpiration under conditions of WD+HS combination (Figure 2), is that this process helps to lower the inner flower temperature. This could be highly important for protecting the reproductive processes occurring within the flowers of pseudocleistogamous plants such as soybean. To test whether reducing flower transpiration, by sealing stomatal apertures will cause an increase in inner flower temperature under conditions of WD+HS, we used a thin layer of wax to cover flowers (Stage II and III) of plants (R2 stage) grown under CT, WD, HS and WD+HS conditions, and measured their inner flower temperature. As shown in Figure 4B, sealing stomatal pores with a thin wax layer caused a significant increase of 2-3°C in inner flower temperature of flowers grown under CT, HS or WD+HS conditions. In contrast, the inner flower temperature of flowers from plants subjected to WD did not increase, as the stomata of these flowers were closed. These findings demonstrate that the opening of stomata on sepals of flowers from plants subjected to HS or WD+HS plays an important role in modulating the internal temperature of flowers; potentially mitigating some of the high temperature-derived negative consequences for plant fertilization in cleistogamous plants (Rollins et al., 2013; Mahrookashani et al., 2017; Lawas et al., 2018; Xie et al., 2018; Cohen et al., 2021b; Cohen et al., 2021a; Sinha et al., 2021).

### RNA-Seq analysis of flower buds subjected to a combination of water deficit and heat stress

To gain a better understanding of the different processes occurring within flowers during WD+HS and to compare them to the processes that occur in leaves (Cohen et al., 2021a), we conducted an RNA-Seq analysis of whole flowers (R2, stage II and III; Figure 1B) collected from plants grown under conditions of CT, WD, HS, or WD+HS (Supplemental Datasets S1-S6). Because conditions of WD, HS and WD+HS are likely to affect global processes in all tissues and cell types found in flowers, we did not dissect the flower buds into different tissues. This also allowed us to compare the RNA-Seq data obtained in the current study with a previous RNA-Seq analysis of whole leaves (that also contain multiple tissues and cell types subjected to same conditions; reanalyzed using the same pipeline as described here; Supplemental Datasets S7-S12), performed in the same growth chambers on plants from the same seed batch, under the same growth conditions (Cohen et al., 2021a). As shown in Figure 5A, RT-qPCR analysis conducted on RNA samples, before RNA-Seq analysis, revealed that flowers from plants subjected to the different treatments responded differently. Transcripts encoding cytosolic ascorbate peroxidase 1 (*APX1*), a key ROS metabolizing and signaling enzyme (Davletova et al., 2005; Koussevitzky et al., 2008) primarily accumulated for example in response the HS, while transcripts encoding the key transcriptional regulator dehydration responsive element binding (DREB; Agarwal et al., 2017)-1H primarily accumulated during WD, and transcripts encoding *DREB-1B* primarily accumulated during HS and WD+HS.

**Figure 5.**
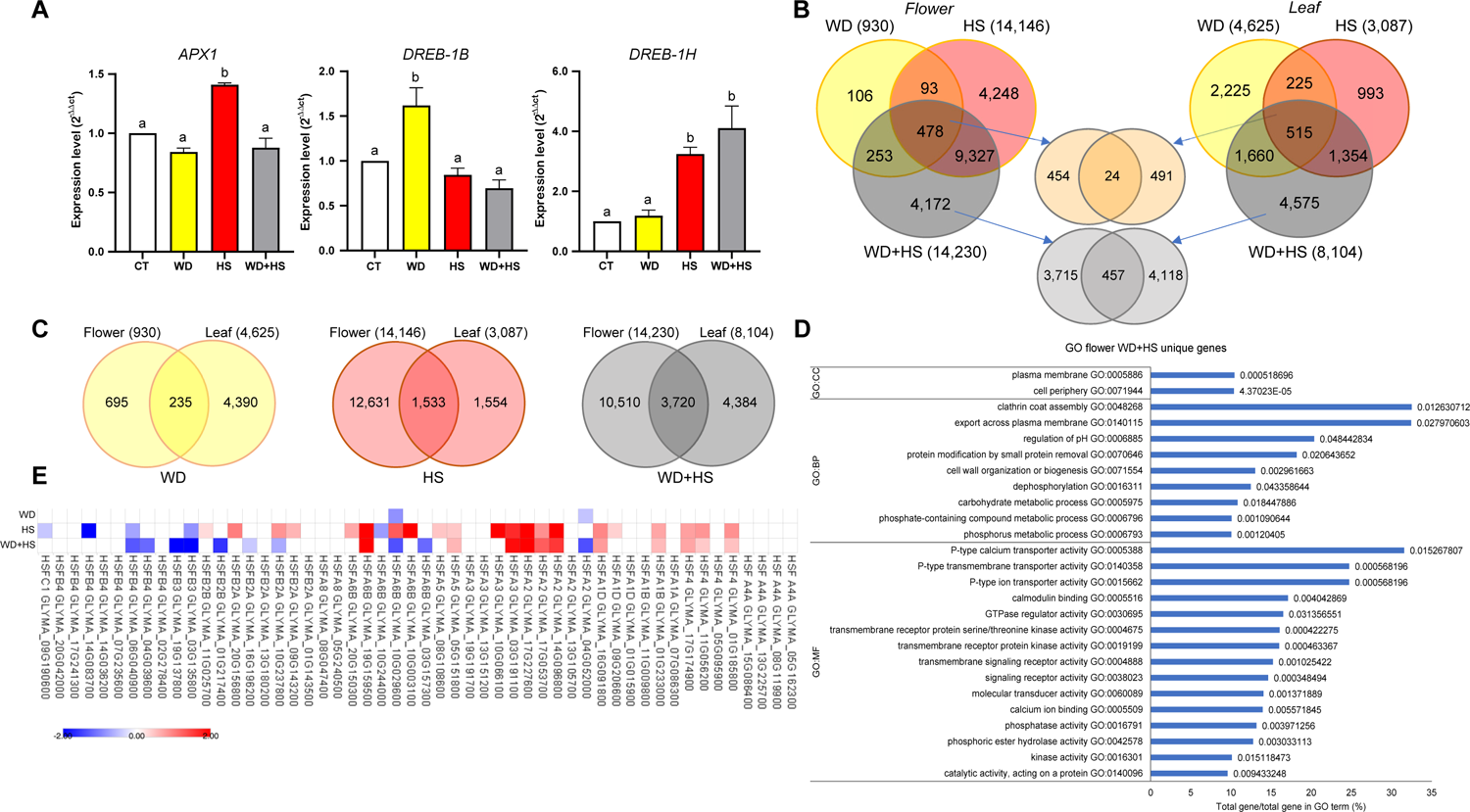
RNA-Seq analysis of flowers subjected to a combination of water deficit and heat stress. **A,** RT-qPCR analysis of ascorbate peroxidase 1 (*APX1*), and dehydration responsive element binding (*DREB*) *1B* and *1H* in flowers from plants subjected to control (CT), heat stress (HS), water deficit (WD), or WD+HS. **B,** Venn diagrams showing the overlap between transcripts with significantly altered expression (up or down regulated) in flowers (left) and leaves (right) in response to HS, WD, or WD+HS. Overlap between transcripts common to all stresses in leaves and flowers is shown in the middle (top) and overlap between transcripts unique to WD+HS in flowers and leaves is shown in the middle (bottom). **C,** Venn diagrams showing the overlap between transcripts with significantly altered expression (up or down regulated) in flower and leaves in response to HS, WD, or WD+HS. **D,** GO enrichment analysis of transcripts unique to WD+HS in flowers (4,177). **E,** Heat map showing the expression pattern of all heat shock transcription factors (*HSFs*) in flowers subjected to HS, WD, or WD+HS. Analysis was performed in 3 biological repeats and all transcripts are significant at P < 0.05 (negative binomial Wald test followed by Benjamini–Hochberg correction). Abbreviations: APX, ascorbate peroxidase; DREB, dehydration responsive element binding; GO, gene ontology; HSF, heat shock transcription factor; CT, control; HS, heat stress; qPCR, quantitative polymerase chain reaction; RT, real-time; WD, water deficit.

Venn diagrams depicting the overlap between transcripts responding to the different treatments in flowers and leaves revealed that in contrast to leaves, flowers accumulated many more transcripts in response to HS (14,146) or WD+HS (14,230), but fewer transcripts in response to WD (930) (Figure 5B; Supplemental Datasets S13-S41). Interestingly, the number of transcripts with a common response to all treatments in flowers (478) and leaves (515) was very similar, suggesting that these transcripts represent a core set of WD-, HS- and WD+HS-response transcripts. However, the overlap between these core sets of leaf and flower transcripts was low (24; Figure 5B) demonstrating that even when it comes to the most common transcripts, the response of flower and leaf tissues to stress is different. A relatively low overlap (457) was also found between transcripts specific for a combination of WD+HS in flowers (4,172) and leaves (4,575) (Figure 5B), further suggesting that the response of flowers to this stress combination is different from that of leaves. A comparison between the overall transcriptomics responses of flowers and leaves to the individual WD, HS, and WD+HS treatments (930, 14,146, and 14,230 in flowers, and 4,625, 3,087, and 8,104 in leaves, respectively) also revealed that these two tissues responded differently (overlap of 235, 1,533, and 3,720, respectively) (Figure 5C). Although some overlap was found between flowers and leaves, in general there were many more flower-specific transcripts that respond to HS and WD+HS (12,613 and 10,510, respectively), and many more leaf specific transcripts that responded to WD (4,390). Overall, the results presented in Figures 5B and C demonstrate that the response of soybean flowers is very different from that of leaves to WD+HS.

Gene ontology (GO) annotation analysis of transcripts with a unique response to WD+HS in flowers (4,172; Figure 5D) revealed that this group of transcripts is enriched in calcium signaling, kinase and protease activity, clathrin-associated vesicle transport, and other types of membrane transport mechanisms and pumps. Because different transcription factor (TF) families, such as heat shock transcription factors (HSFs), MYBs and AP2-EREBP play a critical role in plant acclimation to stress combination (Zandalinas et al., 2020b), we compared the pattern of their expression between leaves and flowers of soybean plants subjected to CT, WD, HS and WD+HS treatments (Supplemental Datasets S42-S44). The pattern of expression of many of these TF families was different between flowers subjected to CT, WD, HS or WD+HS treatments (Supplemental Datasets S42-S44). The pattern of expression of HSFs was for example different between flowers subjected to WD+HS, HS, or WD (Figure 5E). This finding suggest that different types of heat and other stress responses might be activated in flowers when WD and HS are combined (*i.e.,* WD+HS).

Interestingly, the expression of several transcripts encoding the key ABA biosynthetic enzymes zeaxanthin epoxidase (*ABA1*) and 9-cis-epoxycarotenoid dioxygenase (*NCED*) was primarily elevated in flowers from plants subjected to WD or WD+HS, while the expression of several other key ABA biosynthetic enzymes encoding Xanthoxin dehydrogenase (*ABA2*) and aldehyde oxidase (*AAO*) was primarily elevated in flowers from plants subjected to HS or WD+HS (Supplemental Figure S3). In contrast, the expression of several transcripts encoding the key ABA degradation enzyme ABA 8′-hydroxylase (*CYP707A*) was specifically elevated in flowers subject to HS or WD+HS (Supplemental Figure S3). These findings, coupled with the stomatal closure response of sepals from flowers subjected to CT, HS, or WD+HS to ABA application (Figure 4A), suggest that enhanced degradation of ABA in flowers from plants subjected to HS or WD+HS could keep ABA levels suppressed, and therefore stomata open under conditions of HS and WD+HS (Figure 2).

### Suppressed accumulation of ABA and JA in flowers from plants subjected to heat stress or a combination of water deficit and heat stress

To determine the impact of WD+HS on the level of different plant hormones involved in responses to stress, including stomatal regulation (*e.g.,* ABA; Supplemental Figure S3), we measured the levels of ABA, jasmonic acid (JA), salicylic acid (SA), and auxin (IAA) in flowers and leaves from plants subjected to CT, WD, HS and WD+HS (Figure 6; Supplemental Figure S4). In agreement with our findings that transcripts encoding the ABA biosynthetic enzymes *ABA1* and *NCED* are primarily elevated in flowers in response to WD or WD+HS (Supplemental Figure S3), the level of ABA in flowers from plants subjected to WD or WD+HS was higher than that of CT or HS (Figure 6A). In contrast, and in agreement with our findings that transcripts encoding the ABA degradation enzyme *CYP707A* are specifically elevated in flowers in response to HS or WD+HS (Supplemental Figure S3), the levels of dihydrophaseicacid (DPA), a product of ABA degradation, was significantly elevated only in flowers from plants subjected to HS or WD+HS (Figure 6B). These findings support our RNA-Seq analysis (Supplemental Figure S3) and ABA application study (Figure 4A) and demonstrate that an enhanced process of ABA degradation occurs in flowers from plants subjected to HS or WD+HS. Interestingly, compared to flowers from plants subjected to CT or WD stress, flowers from plants subjected to HS or WD+HS contained significantly lower levels of JA and the active form of JA, JA-Isoleucine (JA-Ile; Figures 6C and D). Because both ABA and JA can induce stomatal closure during stress in plants (Nilson and Assmann, 2007; Zandalinas et al., 2016; Zhu, 2016; Hsu et al., 2021; Markham and Greenham, 2021), our findings that flowers from plants subjected to HS or WD+HS contained lower levels of JA (Figure 6C) and JA-Ile (Figure 6D), as well as actively degrade ABA (Figure 6B), provide a hormone-based mechanistic understanding to the opening of stomata on flowers during HS and WD+HS (Figure 2). In contrast to flowers (Figures 6A-D), the levels ABA, JA, and JA-Ile in leaves subjected WD+HS were not suppressed (Figures 6E-G), and stomata on leaves of flowers subjected to WD+HS were closed (Figure 2). Interestingly, compared to leaves from CT or WD stress, the level of IAA was higher in leaves subjected to HS or WD+HS (Supplemental Figure S4). In addition, compared to flowers from plants subjected to WD or WD+HS the level of SA was higher in flowers subjected to HS (Supplemental Figure S4). Further studies are needed to determine the roles of SA and IAA in plant responses to WD+HS.

**Figure 6.**
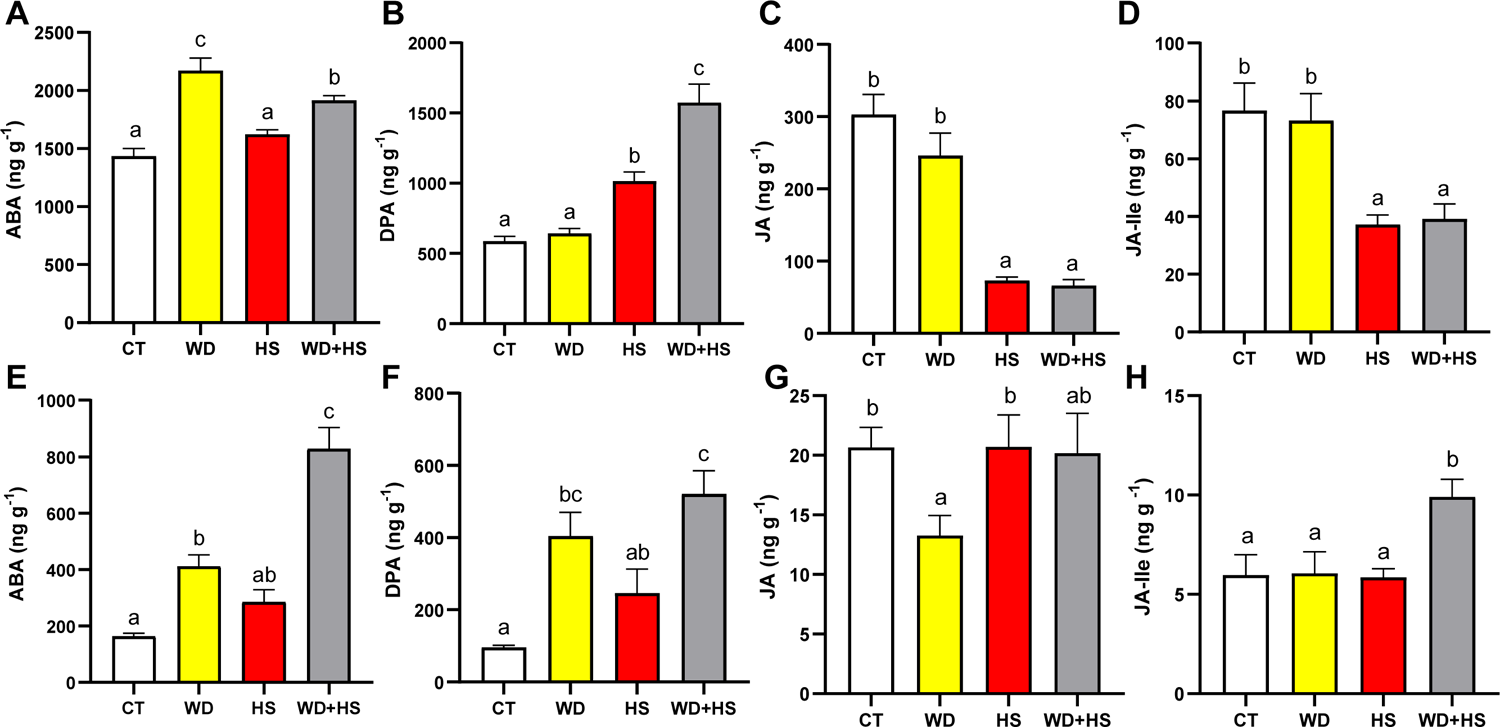
Suppressed accumulation of ABA and JA in flowers from plants subjected to heat stress or a combination of water deficit and heat stress. Levels of ABA (**A**), the ABA degradation product DPA (**B**), JA (**C**), and JA-Ile (**D**) in flowers from plants subjected to control (CT), water deficit (WD), heat stress (HS), or WD+HS. **E-F,** Same as **A-D**, but for leaves. All experiments were repeated at least three times with at least 10 plants per biological repeat. Data are presented as mean ± SE. Different letters denote significance at P < 0.05 (ANOVA followed by a Fisher’s post hoc test). Abbreviations: ABA, abscisic acid; CT, control; DPA, dihydrophaseicacid; HS, heat stress; JA, jasmonic acid; JA-Ile, JA-Isoleucine; WD, water deficit.

### The effect of water deficit and heat stress combination on flower and leaf stomatal aperture, transpiration, and temperature in tobacco

To determine whether stomata of sepals and leaves belonging to a different plant species respond in a similar manner to soybean (Figure 2), we studied the response of *Nicotiana tabacum* (cv SR1, *petite Havana*) plants to WD, HS, and WD+HS. As shown in Figure 7A, and in agreement with our previous analysis of tobacco plants subjected to a combination of WD+HS (Rizhsky et al., 2002), the leaf temperature of plants subjected to a combination of WD+HS was higher than that of plants subjected to WD or HS. This increase was accompanied by closure of stomata and suppressed transpiration. In contrast to leaves, and similar to soybean (Figure 2), stomata on sepals of tobacco plants subjected to WD+HS were open, allowing transpiration to occur (Figure 7B). Although stomata were open and transpiration occurred, the inner flower temperature of tobacco plants subjected to WD+HS (measured for unopened flowers), was higher than that of flowers subjected to HS or WD (Figure 7B; similar to our findings with soybean; Figure 2). As in soybean, it is possible that due to differences in tissue thickness between leaves and flowers, keeping transpiration ongoing in flowers is not sufficient to reduce the inner temperature of flowers more extensively.

**Figure 7.**
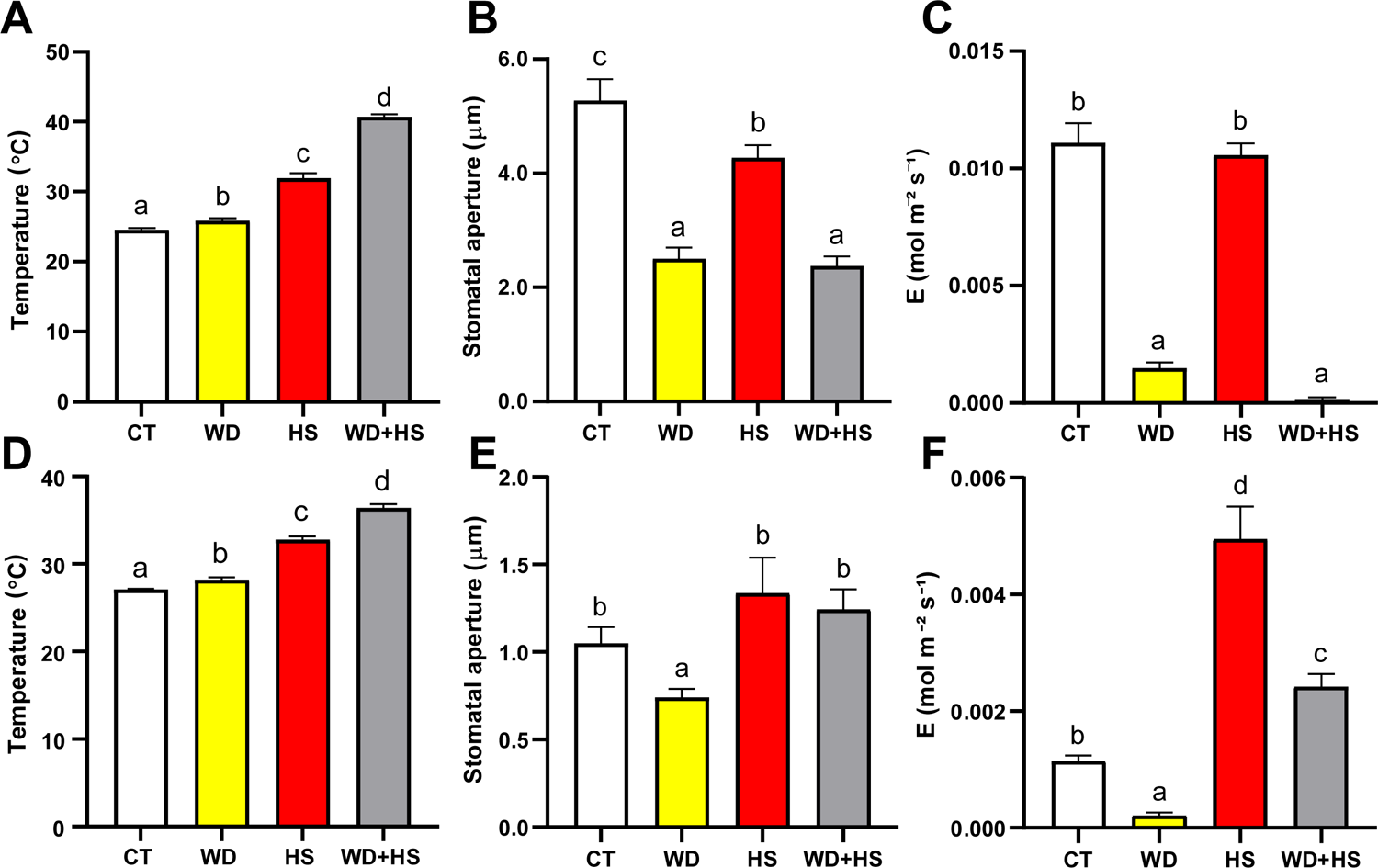
The effect of water deficit and heat stress combination on flower and leaf stomatal aperture, transpiration, and temperature in tobacco. **A,** Leaf temperature of tobacco plants subjected to control (CT), heat stress (HS), water deficit (WD), or WD+HS. **B,** Stomatal aperture of tobacco leaves subjected to CT, WD, HS, or WD+HS. **C,** Transpiration of tobacco leaves subjected to CT, WD, HS, or WD+HS. **D-F,** Same as **A-C**, but for tobacco flowers from plants subjected to CT, WD, HS, or WD+HS. All experiments were repeated at least three times with at least 10 plants per biological repeat. Data are presented as mean ± SE. Different letters denote significance at P < 0.05 (ANOVA followed by a Fisher’s post hoc test). Abbreviations: CT, control; E, transpiration; HS, heat stress; WD, water deficit.

### Yield of soybean and tobacco subjected to a combination of water deficit and heat stress

Our findings that stomata of sepals are open during HS and WD+HS, and that this limits increases in internal flower temperature (Figures 2 and 4), suggest that the opening of stomata on sepals could curb the extent of yield losses that may otherwise be caused by WD+HS. Because the temperature of flowers from plants grown under conditions of HS and WD+HS was comparable (albeit higher in plants subjected to WD+HS; Figures 1, 4 and 7), we hypothesized that yield penalty in plants subjected to WD+HS will be comparable to that of plants subjected to HS alone. To test this hypothesis, we grew soybean and tobacco plants under conditions of CT, WD HS, and WD+HS and scored them for number of flowers, number of pods and seed weight per plant. In contrast to our previous analysis of soybean yield under these conditions (Cohen et al., 2021a), plants were scored for the different parameters while in the chambers, and not following a recovery period in the greenhouse. As shown in Figure 8A, soybean and tobacco plants subjected to HS produced more flowers compared to plants subjected to CT, WD, or WD+HS. The number of pods and seeds produced by plants subjected to HS was however lower than that of plant subjected to CT or WD conditions, suggesting that most of these flowers could not produce pods and seeds (Figures 8B and C). Interestingly, the number of pods and seeds produced in soybean plants subjected to HS or WD+HS was comparable (Figures 8B and C), suggesting that the differential transpiration response of soybean plants (Figures 2 and 4) could help protect flowers during WD+HS. In contrast, HS and WD+HS had a much more severe impact on pod and seed production in tobacco, with WD+HS being the more severe of the two (Figure 8B). Our findings suggest that at least in soybean, that uses pseudocleistogamy for plant reproduction (Takahashi et al., 2001; Khan et al., 2008; Benitez et al., 2010), the differential transpiration of sepals during a combination WD+HS could keep flower temperature under control and help prevent excessive yield losses.

**Figure 8.**
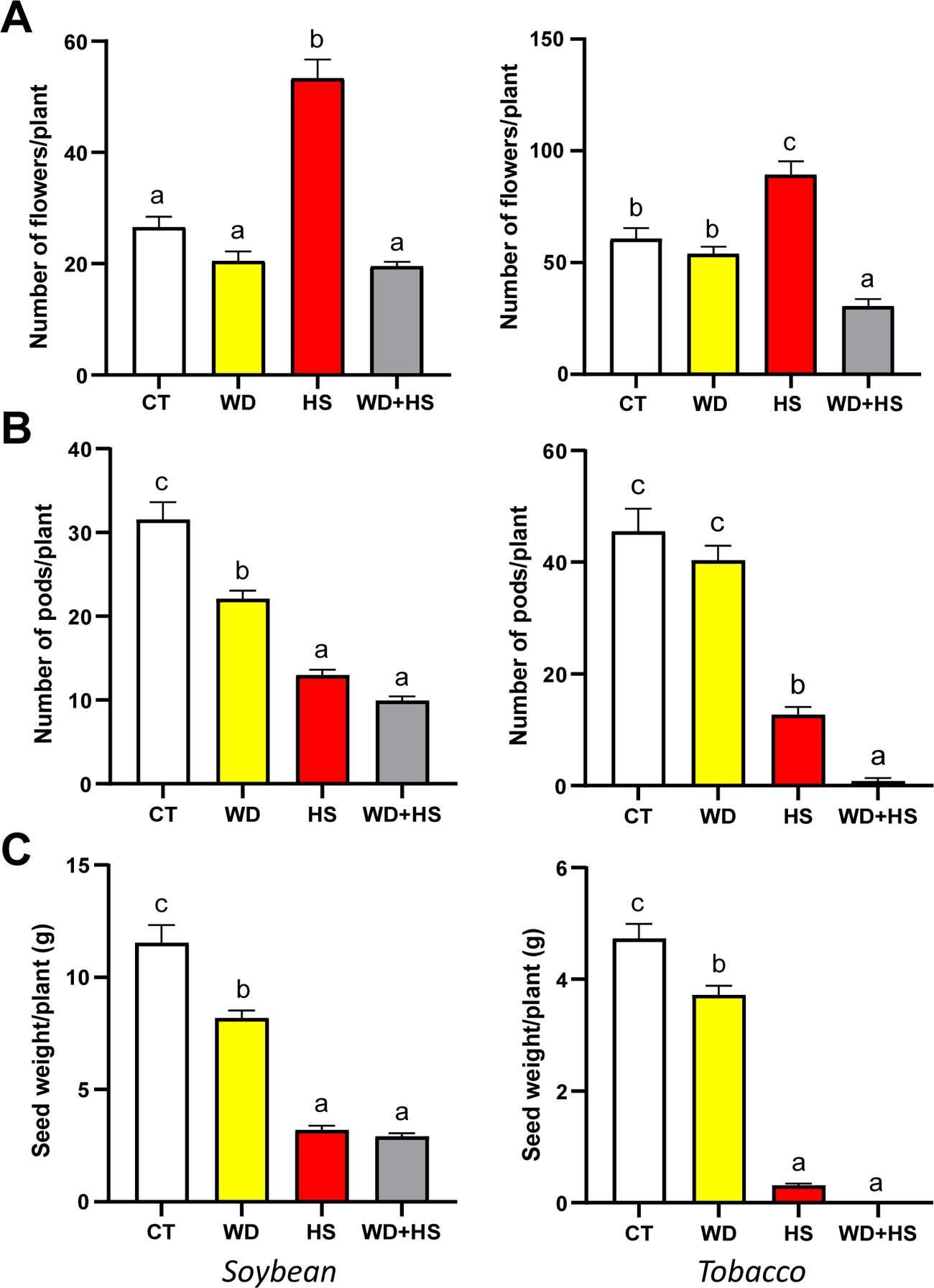
Yield of soybean and tobacco plants subjected to a combination of water deficit and heat stress. **A-C,** Number of flowers per plant (**A**), number of pods per plant (**B**), and total seed weight per plant (**C**), of soybean (left bar graphs) and tobacco (right bar graphs) plants subjected to control (CT), heat stress (HS), water deficit (WD), or WD+HS. All experiments were repeated at least three times with at least 10 plants per biological repeat. Data are presented as mean ± SE. Different letters denote significance at P < 0.05 (ANOVA followed by a Fisher’s post hoc test). Abbreviations: CT, control; HS, heat stress; WD, water deficit.

## Discussion

Heat waves occurring during periods of drought can inflict heavy losses to agricultural production, especially if they occur during the reproductive growth phase of crops (Mittler, 2006; Rollins et al., 2013; Mazdiyasni and AghaKouchak, 2015; Mahrookashani et al., 2017; Lawas et al., 2018; Liu et al., 2020; Cohen et al., 2021b; Rivero et al., 2021; Sinha et al., 2021). Because water is needed to cool the plant via transpiration, we reasoned that when WD is combined with HS it would limit the ability of plants to cool their flowers and cause a severe heat-induced reduction in yield. Here, we show that conditions of WD+HS, that were found to reduce yield in many different crops (Mittler, 2006; Rollins et al., 2013; Mahrookashani et al., 2017; Lawas et al., 2018; Liu et al., 2020; Rivero et al., 2021; Sinha et al., 2021), are indeed accompanied by higher inner flower temperatures (Figures 1 and 6). Higher leaf temperatures were previously reported for plants subjected to WD+HS and linked to the closure of stomata on leaves during stress combination (Rizhsky et al., 2002; Rizhsky et al., 2004; Carmo-Silva et al., 2012; Zandalinas et al., 2016; Cohen et al., 2021a). We therefore expected stomata of flowers from plants subjected to WD+HS to also be closed. Surprisingly, however, they were open (Figures 2 and 7). Moreover, transpiration rates of flowers from plants subjected to WD+HS were as high as those of flowers subjected to HS alone (Figures 2 and 7). In contrast to flowers, stomata on leaves of plants subjected to WD+HS were closed (Figures 2 and 7). Our results therefore reveal that during a combination of WD+HS plants prioritize transpiration through flowers over transpiration through leaves by opening their sepal stomata, while keeping their leaf stomata closed (Figure 9). This ‘differential transpiration’ strategy lowers flower internal temperature (Figure 4) and enables some reproductive processes to occur (Figure 8). Under conditions of WD+HS the plant might therefore attempt to protect reproductive processes, at the expense of vegetative organs. This acclimation strategy could also prove effective in other scenarios that may result in higher inner flower temperatures (*e.g.,* combinations of pathogen infection, mechanical wounding, high CO_2_, or air pollution, such as ozone, that cause stomal closure with heat stress; Melotto et al., 2006; Vahisalu et al., 2010; Raven, 2014; Deger et al., 2015; Chen et al., 2017; Zhang et al., 2018; Devireddy et al., 2020).

**Figure 9.**
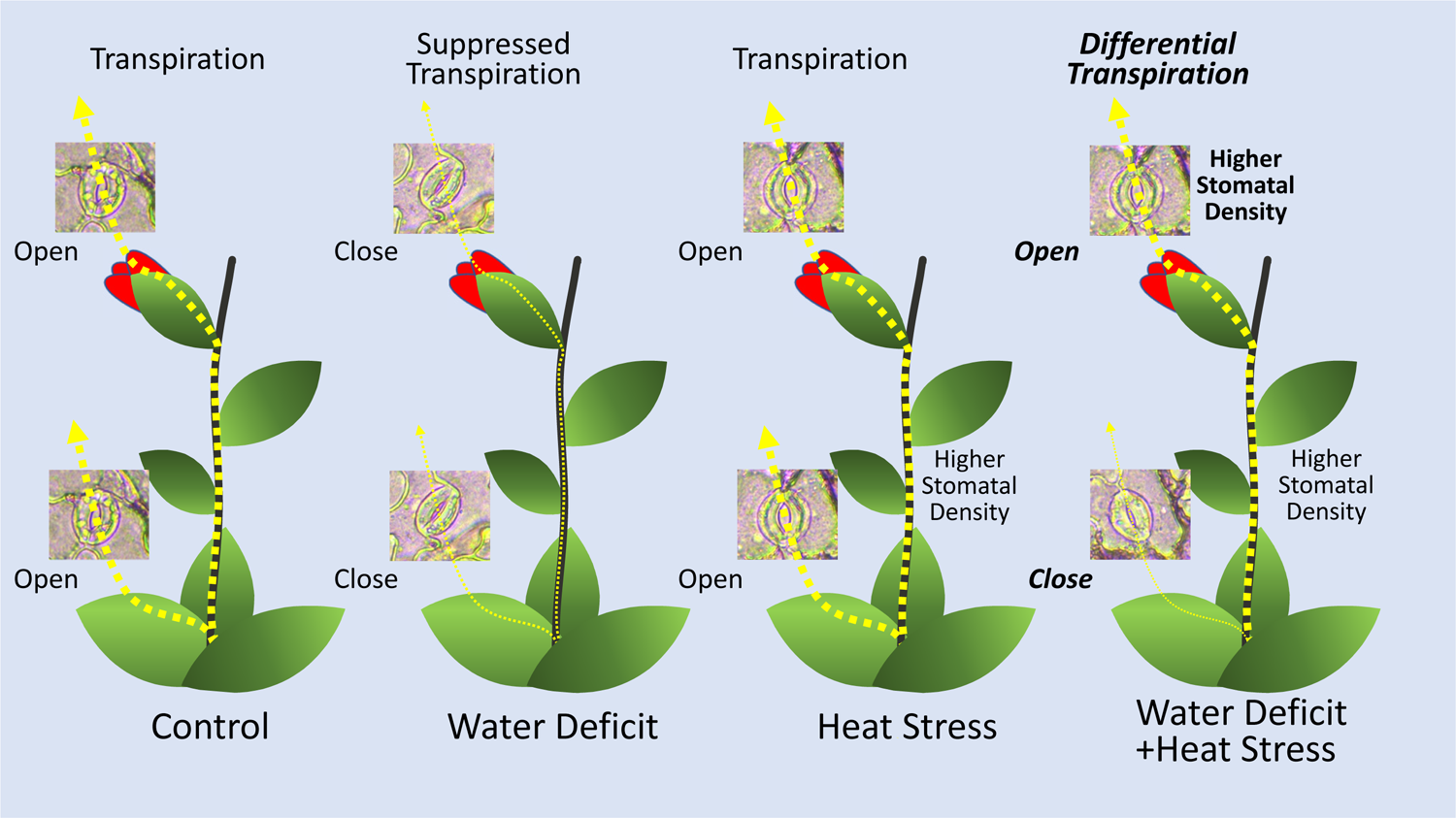
A model depicting ‘differential transpiration’ during a combination of water deficit and heat stress. Control plants (left) conduct transpiration through open stomata on their leaves and flowers. In response to water deficit, plants (second from left) close their stomata on leaves and flowers and suppress transpiration. During heat stress, plants (third from left) keep their stomata on leaves and flowers open and maintain transpiration. In contrast, during a combination of water deficit and heat stress, plants (right) keep their stomata on flowers open, while closing their stomata on leaves. This strategy of differential transpiration allows plants subjected to the stress combination to cool their flowers and limit heat-induced negative impacts on yield. In addition, the stomatal density of flowers from plants subjected to a combination of water deficit and heat stress is higher.

Enhanced transpiration of flowers grown under conditions of WD+HS could cause flowers to undergo desiccation due to limited water resources. Indeed, flowers of plants subjected to WD+HS had a lower water potential (Ψ) compared to flowers of plants subjected to HS alone (Figure 2). This observation suggests that the strategy of differential transpiration under conditions of WD+HS (Figure 9) has its limits, and once flowers will reach a desiccation point of no return, reproductive processes will be further, and perhaps irreversibly, damaged. The challenges faced by flowers of plants subjected to WD+HS, and the findings that they may be subjected to a combination of heat and desiccation stress (Figure 2), is also reflected in our RNA-Seq analysis. Flowers from plants subjected to WD+HS displayed a unique transcriptomics response that was different than that of flowers from plants subjected to HS or WD (Figures 5A and B). Interestingly, the overall transcriptomics response of flowers to WD, HS, or WD+HS was different from that of leaves (with the highest degree of similarity observed between flowers and leaves subjected to a combination of WD+HS; Figure 5C). This observation might reflect the many different reproductive processes that occur in developing flowers compared to leaves but could also suggest that flowers are subjected to different types or degrees of stress compared to leaves under conditions of WD+HS. Further studies are of course needed to address these intriguing possibilities.

Our RNA-Seq analysis further revealed that the expression of several transcripts encoding the key ABA degradation enzymes ABA 8′-hydroxylase is specifically enhanced in flowers from plants subjected to HS, or WD+HS (Supplemental Figure S3). Interestingly, stomata of flowers grown under HS, or WD+HS closed in response to external application of ABA (Figure 4), suggesting that ABA may not accumulate to high levels in flowers from plants subjected to these stresses. ABA biosynthesis might therefore occur in flowers from plants subjected to WD, HS, or WD+HS, but under conditions of HS and WD+HS ABA degradation could keep ABA levels low and therefore stomata open (Figure 2). Indeed, the levels of the ABA degradation product DPA were specifically and significantly elevated in flowers from plant subjected to HS or WD+HS (Figure 6B), supporting this hypothesis. In addition to enhanced ABA degradation (Figure 6B; Supplemental Figure S3), the levels of JA and JA-Ile were also specifically and significantly altered (reduced) in flowers from plants subjected to HS or WD+HS (Figures 6C and D). Because JA-Ile and ABA are both involved in the regulation of stomatal closure during stress (Nilson and Assmann, 2007; Zandalinas et al., 2016; Zhu, 2016; Hsu et al., 2021; Markham and Greenham, 2021; Zhang et al., 2021), the acclimation strategy of ‘differential transpiration’ during WD+HS (Figure 9) could be explained by differential accumulation of these two hormones between flowers (enhanced degradation of ABA and reduced levels of JA-Ile; Figures 6B and D), and leaves (enhanced accumulation of ABA and JA-Ile; Figures 6A and D). Further studies are of course needed to determine how genes involved in the biosynthesis, degradation, and transport of these two hormones are differentially regulated in flowers and leaves in response to WD, HS, WD+HS, and other stressful conditions.

For plants that use cleistogamy or pseudocleistogamy for reproduction, cooling of flowers by opening stomata on sepals could be especially important to protect reproductive processes from high temperatures. Although most of the closed soybean flower surface is covered by sepals (Figure 1B), cooling of a closed flower that has a diameter of about 1.5-2 mm by transpiration from sepals is much harder than cooling a leaf that has a thickness of about 0.12-0.15 mm. In addition, although sepal stomata are bigger than leaf stomata (Supplemental Figure S2), their density is lower (Figure 3). It is likely that due to the thickness of flower buds, cooling by transpiration can only contribute to a reduction of about 2-3°C in flower temperature during conditions of HS or WD+HS (Figure 4) and this reduction would of course depend on the water status of the plant. It therefore seems logical to speculate that the smaller the closed flower is, the easier it will be to cool it by transpiration. Moreover, the strategy of differential transpiration, revealed by this work (Figure 9), may be primarily beneficial for annual plants that need to produce seeds every season, as opposed to perennials that need to protect their vegetative tissues and might abort or skip flowering during entire seasons if conditions are not permissive. Because many important crops, such as soybean, wheat, and barely are annual, use cleistogamy or pseudocleistogamy for reproduction, and have relatively small flowers, the strategy of differential transpiration could play an important economic role in preventing yield penalty under different stress conditions, especially when they occur during the reproductive stage of plant growth. Further studies are of course needed to dissect the different pathways involved in this response and identify key regulators that control it.

The identification of differential transpiration as a potential mechanism that prevents yield losses under conditions of WD+HS combination highlights new avenues for crop improvements. For example, the density and size of stomata on sepals or other floral organs might be altered to improve transpirational cooling of reproductive tissues. In addition, the pathways and mechanisms controlling stomatal responses of flowers could be manipulated to modulate opening or closing, depending on different environmental conditions, protecting flowers from overheating. These manipulations could target the timing of opening or closing as well as the different stimuli and stresses that trigger them.

In summary, we report a novel acclimation strategy of plants that prioritize the transpiration of reproductive tissues over that of vegetative tissues (Figure 9). This mechanism, termed ‘differential transpiration’, protects flowers of plants from overheating and could be important to minimize yield losses under conditions of stress combination. In addition, it can serve as a new example for plant plasticity in responses to abiotic stress.

## MATERIALS AND METHODS

### Plant material and stress treatments

Soybean (*Glycine max*, cv *Magellan*) seeds were inoculated with *Bradyrhizobium japonicum* inoculum (N-DURE, Verdesian Life Sciences, NC, USA) and germinated in Promix HP (Premier Tech Horticulture; PA, USA) under short day growth condition (12-h light/12-h dark), 500 μmol photons m^-2^ s^-1^, at 28/24 ℃ day/night temperature (the temperature was linearly ramped from 24 to 28 °C between 6.00-8.00 AM and linearly decreased to 24 °C from 16.00-20.00 PM), for a week in growth chambers (BDR16, Conviron; Canada), as previously described (Cohen et al., 2021a). After a week, seedlings from trays were transplanted into pots containing 1 kg of Promix HP soaked in 1 l of water-fertilizer (Zack’s classic blossom booster 10-30-20; JR peters Inc., USA) mix (Cohen et al., 2021a). Plants were then grown for the next 16-18 days (until start of first open flower, R1 developmental stage, Fehr et al., 1971) under 28/24 ℃ day/night temperatures as above, but the light intensity (12-h light/12-h dark photoperiod) was increased to 1000 μmol photons m^-2^ s^-1^. Plants were fertilized twice a week (Cohen et al., 2021a). At R1 plants in the WD and WD+HS treatments were supplied with 30% of the water available for transpiration (determined by weighing pots daily as described in Cohen et al., 2021a), while plants in the CT and HS treatments were well watered. Plants in the HS and WD+HS treatments were further subjected to HS by ramping the temperature from 28 to 38 °C between 6.00-8.00 AM and decreasing it down to 28 °C between 16.00-20.00 PM. All measurements were conducted 10 days following the start of the stress treatments using new flowers and leaves that developed under the stress conditions (R2 stage).

### Temperature, gas exchange and water potential

Flower temperature was measured with a microthermocouple sensor (Physitemp instruments LLC; Clifton, NJ, USA) by inserting the hypodermal needle microprobe (Physitemp instruments LLC; Clifton, NJ, USA) 0.75-1 mm into soybean flowers and 1.5-2 mm into tobacco flowers. Data was recorded using a Multi-Channel Thermocouple Temperature Data Logger (TCTemp X-Series, ThermoWorks LogMaster; UT, USA) between 11.30 AM-12:30 PM. Stomatal conductance, transpiration, leaf temperature, and photosynthesis were recorded using a LICOR Portable Photosynthesis System (LI-6800, LICOR, Lincoln, NE, USA) between 12.00-1.00 PM as described previously (Cohen et al., 2021a). Leaf temperature was also recorded using infrared camera (FLIR C2, FLIR systems AB; Wilsonville, OR, USA) as previously described (Zandalinas et al., 2020a). Water potential of leaf discs (8 mm) and flowers (cut longitudinally into half) from plants was measured using Dewpoint Potentiameter (WP4C, METER Group, Inc. WA, USA) as described previously (Cohen et al., 2021a). Water potential (leaf and flowers) was measured from 5-6 plants (from each treatment) between 12.00-4.00 PM.

### Measurements of stomatal aperture and stomatal density

Adaxial and abaxial surface of leaves and outer surface of sepals from soybean and tobacco plants were pasted onto microspore slide, between 11.00 AM-12.00 PM, with a medical adhesive (Hollister Adapt 7730; Libertyville, IL, USA), as previously described (Devireddy et al., 2020; Zandalinas et al., 2020a). Stomatal aperture measurements were performed using an EVOS XL microscope (Invitrogen by Thermo Fisher Scientific, CA, USA), as described previously (Devireddy et al., 2020; Zandalinas et al., 2020a). Both width and length of stomatal aperture were measured using ImageJ (https://imagej.nih.gov/ij). Number of epidermal cells and stomata per microscopic field of view were counted to calculate stomatal density. Flowers at different stages were also fixed in 4% paraformaldehyde, mounted in paraffin, sectioned, and stained at the Histochemistry Diagnostic Laboratory at the University of Missouri, Columbia.

### Yield measurements

Yield and reproductive traits were measured as described in (Cohen et al., 2021a), except that plants were scored while still growing inside the chambers, and not following recovery under greenhouse conditions. Number of flowers and pods were counted from 10-15 different plants per treatment, for both soybean and tobacco, and represented as number of flower/pods per plant. Seeds from each plant (10-15 different plants) were pooled and weighed for both soybean and tobacco.

### ABA and wax application

ABA (50 µM, Sigma-Aldrich, St. Louis, MO, USA) was sprayed on flowers of soybean plants (R2), growing under the different stress conditions, as previously described (Zandalinas et al., 2016). Control flowers were sprayed with water. Plants were then returned to the chambers and stomatal aperture was measured 60 min post ABA application. To seal stomata, Vaseline® (Sigma-Aldrich, St. Louis, MO, USA) was gently applied to flowers of plants growing under the different stress conditions using Q-tips. Plants were then returned to the chambers and flower temperature was recorded as described above 120 min post wax application.

### Hormone measurements

Hormone extraction and analysis were performed as described in (Zandalinas et al., 2016; Balfagón et al., 2019). Briefly, 10 mg of dry tissue was extracted in 2 mL of ultrapure water after spiking with 50 ng of [d_6_]-ABA, [C_13_]-SA and dihydrojasmonic acid, and 5 ng of [d_2_]-IAA in a ball mill (MillMix20, Domel, Železniki, Slovenija). After centrifugation at 10,000 g at 4°C for 10 min, the supernatant was recovered, and pH adjusted to 3 with 30% acetic acid. The water extract was partitioned twice against 2 mL of diethyl ether and the organic layer recovered and evaporated under vacuum in a centrifuge concentrator (Speed Vac, Jouan, Saint Herblain Cedex, France). Once dried, the residue was resuspended in a 10:90 MeOH:H_2_O solution by sonication. The resulting solution was filtered through 0.22 µm polytetrafluoroethylene membrane syringe filters (Albet S.A., Barcelona, Spain) and directly injected into an ultraperformance LC system (Acquity SDS, Waters Corp., Milford, MA, USA). Chromatographic separation was conducted on a reversed-phase C18 column (Gravity, 50 × 2.1 mm, 1.8-µm particle size, Macherey-Nagel GmbH, Germany) using a MeOH:H_2_O (both supplemented with 0.1% acetic acid) gradient at a flow rate of 300 µL min^−1^. Hormones were quantified with a TQS triple quadrupole mass spectrometer (Micromass, Manchester, UK) connected online to the output of the column though an orthogonal Z-spray electrospray ion source. All data were acquired and processed using Mass Lynx v4.1 software.

### RNA isolation and RT-qPCR

Soybean flowers (Stage II and III, from R2 plants) were collected from plants between 11.30 AM-12:30 PM and immediately frozen in liquid nitrogen. About 30-40 flowers were pooled together from 8-10 different plants for each biological repeat and RNA was isolated using RNAeasy plant mini kit (Qiagen, Germantown, MD, USA). RNA was converted to cDNA using PrimeScript RT Master Mix (Takara). Real-time quantitative PCR (RT-qPCR) was performed with gene-specific primers (Supplementary Table 1) using EF1α as internal reference using the CFX Connect Real-Time PCR Detection System (Bio-Rad, Hercules, CA, USA) as previously described in (Zandalinas and Mittler, 2021).

### RNA sequencing and data analysis

RNA libraries for sequencing were prepared using standard Illumina protocols and RNA sequencing was performed by Novogene co. Ltd (https://en.novogene.com/; Sacramento, CA, USA) using NovaSeq 6000 PE150. Read quality control was performed using Trim Galore v0.6.4 (https://www.bioinformatics.babraham.ac.uk/projects/trim_galore/) & FastQC v0.11.9 (https://www.bioinformatics. babraham.ac.uk/projects/fastqc/). The RNA-seq reads were aligned to the reference genome for Soybean - Glycine max v2.1 (downloaded from ftp://ftp.ensemblgenomes.org/pub/plants/release-51/fasta/glycine_max/dna/), using Hisat2 short read aligner (Kim et al., 2019). Intermediate file processing of sam to sorted bam conversion was carried out using samtools v1.9 (Danecek et al., 2021). Transcript abundance in levels expressed as FPKM was generated using the Cufflinks tool from the Tuxedo suite (Trapnell et al., 2012) guided by genome annotation files downloaded from the same source. Differential gene expression analysis was performed using Cuffdiff tool (Trapnell et al., 2013), also from the same Tuxedo suite. Differentially expressed transcripts were defined as those that had a fold change with an adjusted P < 0.05 (negative binomial Wald test followed by Benjamini–Hochberg correction). Functional annotation and quantification of overrepresented GO terms (P value < 0.05) and KEGG pathway enrichment (P value < 0.05) were conducted using g:profiler (Raudvere et al., 2019). Venn diagrams were created in VENNY 2.1 (BioinfoGP, CNB-CSIC). Venn diagram overlaps were subjected to hypergeometric testing using the R package phyper (Zandalinas et al., 2020a). Heatmaps were generated in Morpheus (https://software.broadinstitute.org/morpheus).

## Statistical Analysis

All experiments were repeated at least 3 times, each with at least 3 technical repeats. Data is represented as mean ± SE. Statistical analysis was performed using one-way ANOVA followed by Tukey’s or Fisher’s post hoc test (P < 0.05) in GraphPad. Different letters denote statistical significance at P < 0.05.

## SUPPLEMENTAL DATA

**Supplemental Figure S1.**
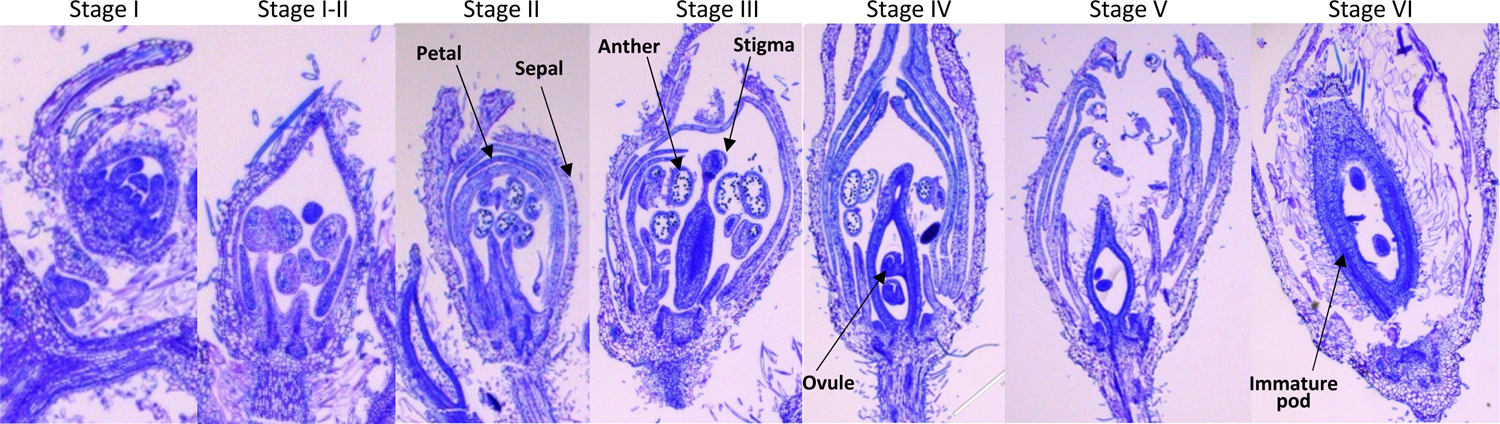
Cross-section light microscopy analysis of fixed and embedded soybean flowers from plants grown under controlled growth conditions.

**Supplemental Figure S2.**
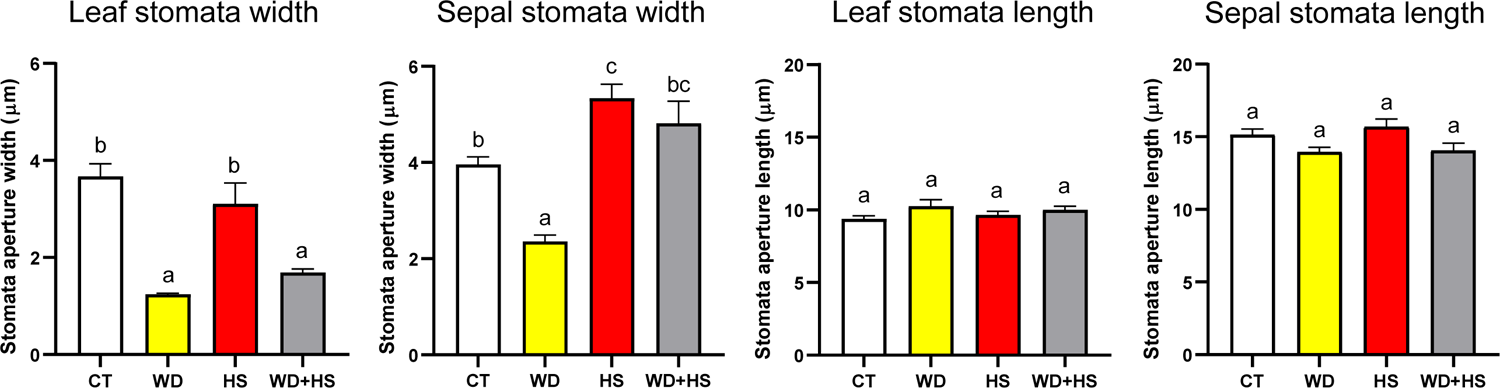
Stomatal width and length of leaves and sepals from plants grown under control (CT), water deficit (WD), heat stress (HS), or WD+HS.

**Supplemental Figure S3.**
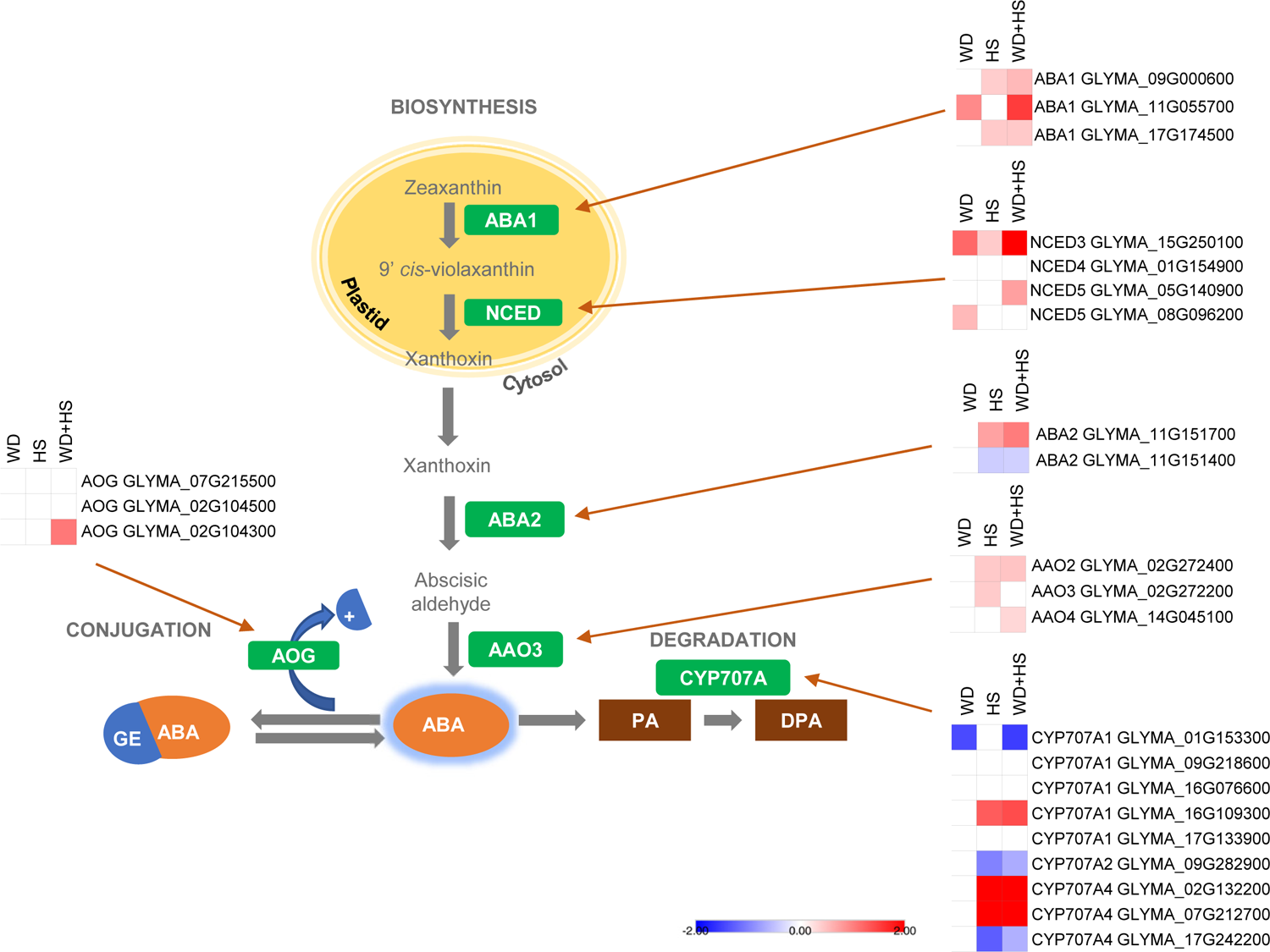
Expression of transcripts involved in abscisic acid (ABA) biosynthesis and degradation in whole flowers from plants grown under control (CT), water deficit (WD), heat stress (HS), or WD+HS.

**Supplemental Figure S4.**
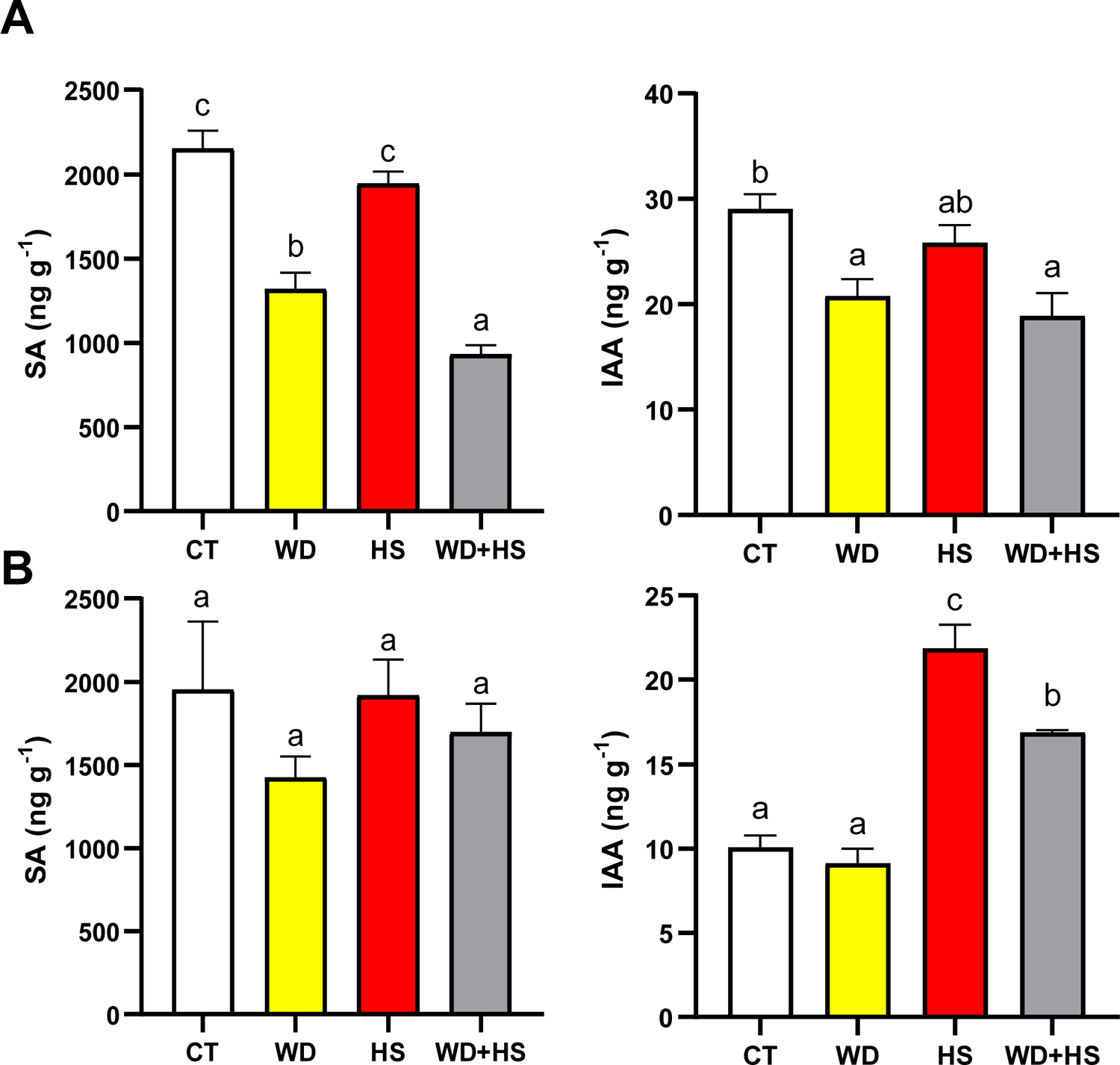
Accumulation of SA and IAA in flowers from plants subjected to heat stress or a combination of water deficit and heat stress.

**Supplemental Table 1.** List of primers used for RT-PCR

**Supplemental Data Set 1.** Transcripts significantly upregulated in soybean flowers subjected to water deficit stress (Figure 5B).

**Supplemental Data Set 2.** Transcripts significantly downregulated in soybean flowers subjected to water deficit stress (Figure 5B).

**Supplemental Data Set 3.** Transcripts significantly upregulated in soybean flowers subjected to heat stress (Figure 5B).

**Supplemental Data Set 4.** Transcripts significantly downregulated in soybean flowers subjected to heat stress (Figure 5B).

**Supplemental Data Set 5.** Transcripts significantly upregulated in soybean flowers subjected to combination of water deficit and heat stress (Figure 5B).

**Supplemental Data Set 6.** Transcripts significantly downregulated in soybean flowers subjected to combination of water deficit and heat stress (Figure 5B).

**Supplemental Data Set 7.** Transcripts significantly upregulated in soybean leaves subjected to water deficit (Figure 5B).

**Supplemental Data Set 8.** Transcripts significantly downregulated in soybean leaves subjected to water deficit (Figure 5B).

**Supplemental Data Set 9.** Transcripts significantly upregulated in soybean leaves subjected to heat stress (Figure 5B).

**Supplemental Data Set 10.** Transcripts significantly downregulated in soybean leaves subjected to heat stress (Figure 5B).

**Supplemental Data Set 11.** Transcripts significantly upregulated in soybean leaves subjected to combination of water deficit and heat stress (Figure 5B).

**Supplemental Data Set 12.** Transcripts significantly downregulated in soybean leaves subjected to combination water deficit and heat stress (Figure 5B).

**Supplemental Data Set 13.** Transcripts exclusively differentially expressed in soybean flowers subjected to water deficit (Figure 5B).

**Supplemental Data Set 14.** Transcripts exclusively differentially expressed in soybean flower subjected to heat stress (Figure 5B).

**Supplemental Data Set 15.** Transcripts exclusively differentially expressed in soybean flower subjected to combination of water deficit and heat stress (Figure 5B).

**Supplemental Data Set 16.** Transcripts commonly expressed in soybean flower subjected to water deficit, and combination of water deficit and heat stress (Figure 5B).

**Supplemental Data Set 17.** Transcripts commonly expressed in soybean flower subjected to water deficit stress and heat stress (Figure 5B).

**Supplemental Data Set 18.** Transcripts commonly expressed in soybean flowers subjected to heat stress, and combination of water deficit and heat stress (Figure 5B).

**Supplemental Data Set 19.** Transcripts commonly expressed in soybean flowers subjected to water deficit, heat stress, and combination of water deficit and heat stress (Figure 5B).

**Supplemental Data Set 20.** Transcripts exclusively expressed in soybean leaves subjected to water deficit (Figure 5B).

**Supplemental Data Set 21.** Transcripts exclusively expressed in soybean leaves subjected to heat stress (Figure 5B).

**Supplemental Data Set 22.** Transcripts exclusively expressed in soybean leaves subjected to combination of water deficit and heat stress (Figure 5B).

**Supplemental Data Set 23.** Transcripts commonly expressed in soybean leaves subjected to water deficit stress and heat stress (Figure 5B).

**Supplemental Data Set 24.** Transcripts commonly expressed in soybean leaves subjected to heat stress, and combination of water deficit and heat stress (Figure 5B).

**Supplemental Data Set 25.** Transcripts commonly expressed in soybean leaves subjected to water deficit, and combination of water deficit and heat stress (Figure 5B).

**Supplemental Data Set 26.** Transcripts commonly expressed in soybean leaves subjected to water deficit, heat stress, and combination of water deficit and heat stress (Figure 5B).

**Supplemental Data Set 27.** Transcripts exclusive to soybean flowers in response to water deficit, heat stress, and combination of water deficit and heat stress compared to leaves (Figure 5B).

**Supplemental Data Set 28.** Transcripts exclusive to soybean leaves in response to water deficit, heat stress, and combination of water deficit and heat stress compared to soybean flowers (Figure 5B).

**Supplemental Data Set 29.** Unique transcripts in response to combination of water deficit and heat stress exclusive to soybean flower compared to leaves (Figure 5B).

**Supplemental Data Set 30.** Unique transcripts in response to combination of water deficit and heat stress exclusive to soybean leaves compared to flower (Figure 5B).

**Supplemental Data Set 31.** Transcripts commonly expressed in soybean flowers and leaves when subjected to water deficit, heat stress, and combination of water deficit and heat stress (Figure 5B).

**Supplemental Data Set 32.** Unique transcripts in response to combination of water deficit and heat stress common between soybean flower and leaves (Figure 5B).

**Supplemental Data Set 33.** Transcripts exclusively expressed in soybean flowers compared to soybean leaves when subjected to water deficit (figure 5C).

**Supplemental Data Set 34.** Transcripts exclusively expressed in soybean leaves compared to soybean flowers when subjected to water deficit (figure 5C).

**Supplemental Data Set 35.** Transcripts commonly expressed in soybean flowers and soybean leaves when subjected to water deficit (figure 5C).

**Supplemental Data Set 36.** Transcripts exclusively expressed in soybean flowers compared to soybean leaves when subjected to heat stress (figure 5C).

**Supplemental Data Set 37.** Transcripts exclusively expressed in soybean leaves compared to soybean flowers when subjected to heat stress (figure 5C).

**Supplemental Data Set 38.** Transcripts commonly expressed in soybean flowers and soybean leaves subjected to heat stress (figure 5C).

**Supplemental Data Set 39.** Transcripts exclusively expressed in soybean flowers compared to soybean leaves subjected to combination of water deficit and heat stress (figure 5C).

**Supplemental Data Set 40.** Transcripts exclusively expressed in soybean leaves compared to soybean flowers subjected to combination of water deficit and heat stress (Figure 5C).

**Supplemental Data Set 41.** Transcripts commonly expressed in soybean flowers and soybean leaves subjected to combination of water deficit and heat stress (Figure 5C).

**Supplemental Data Set 42.** Expression of heat shock factor (HSF) transcripts in soybean flowers and leaves subjected to water deficit, heat stress and combination of water deficit and heat stress (Figure 5E).

**Supplemental Data Set 43.** Expression of MYB transcripts in soybean flowers and leaves subjected to water deficit, heat stress and combination of water deficit and heat stress.

**Supplemental Data Set 44.** Expression of APETALA 2 (AP2) transcripts in soybean flowers and leaves subjected to water deficit, heat stress and combination of water deficit and heat stress.

## Competing interests

The authors declare no competing interests.

## Data Availability

Data supporting the findings of this work are provided in the main paper and Supplemental data. Raw and processed RNA-seq data files were deposited in the GEO database (www.ncbi.nlm.nih.gov/geo/) (accession GSE186317).

## Acknowledgments

This work was supported by funding from the National Science Foundation (IOS-2110017, IOS-1353886, MCB-1936590, IOS-1932639), Interdisciplinary Plant Group, and University of Missouri.

## Author Contributions

R.S., S.I.Z., and Y.F. performed experiments and analyzed the data. S.S. and T.J. analyzed the transcriptomics data, R.M., F.B.F, R.S., T., S.S., A.G.C., S.I.Z. and Y.F. designed experiments, analyzed the data, and/or wrote the manuscript.

## Notes

### Competing Interest Statement

The authors have declared no competing interest.

